# Sexual conflict and STIs: coevolution of sexually antagonistic host traits with a sexually transmitted infection

**DOI:** 10.1101/203695

**Authors:** Alison M. Wardlaw, Aneil F. Agrawal

**Author notes:** Corresponding Author: Alison M. Wardlaw Cargill Building (MPGI) Room 250 1500 Gortner Ave. St. Paul, MN 55108 USA. Contributions: AMW and AFA conceived the idea, designed the model, and wrote the paper. AMW performed the analysis and wrote the simulations. **Data:** Should the manuscript be accepted, Mathematica and Python code will be archived in Dryad and the data DOI will be included in the article.

## Abstract

In many taxa, there is a conflict between the sexes over mating rate. The outcome of sexually antagonistic coevolution depends on the costs of mating and natural selection against sexually antagonistic traits. A sexually transmitted infection (STI) changes the relative strength of these costs. We study the three-way evolutionary interaction between male persistence, female resistance, and STI virulence for two types of STIs: a viability-reducing STI and a reproduction-reducing STI. A viability-reducing STI escalates conflict between the sexes. This leads to increased STI virulence (i.e., full coevolution) if the costs of sexually antagonistic traits occur through viability but not if the costs occur through reproduction. In contrast, a reproduction-reducing STI de-escalates the sexual conflict but STI virulence does not coevolve in response. We also investigated the establishment probability of STIs under different combinations of evolvability. Successful invasion by a viability-reducing STI becomes less likely if hosts (but not parasite) are evolvable, especially if only the female trait can evolve. A reproduction-reducing STI can almost always invade because it does not kill its host. We discuss how the evolution of host and parasite traits in a system with sexual conflict differ from a system with female mate choice.

## Introduction

Sexual conflict over mating rate arises when male reproductive success increases with mating rate while female reproductive success is maximized at some intermediate rate (Bateman, 1948; Arnqvist and Rowe, 2005). Males evolve persistence traits to increase their mating or fertilization rate that often cause harm to females physically or physiologically. Females, in turn, evolve resistance traits that deter males or offset the physiological harm. This conflict gives rise to sexually antagonistic coevolution between male persistence traits and female resistance traits, the outcome of which determines the mating rate. Sexually transmitted infections (STIs) are, by definition, transmitted during mating and thus may play an important role in the evolution of host mating strategies. Here, we explore the interplay between STI virulence evolution and the evolution of host traits mediating sexual conflict over mating rate.

The virulence of STIs that affect host mortality are predicted to evolve in much the same way as that of an ordinary infectious disease, i.e., evolutionary stable virulence is proportional to the natural host mortality rate and depends on the shape of the trade-off between transmission and virulence (Knell, 1999). Surprisingly, evolutionarily stable virulence of an STI does not depend on the mating rate, though its spread and infection prevalence does (Lipsitch and Nowak, 1995; Knell, 1999). STIs, in turn, are known to affect the evolution of mating strategies. Theory has focused on STIs in hosts with conventional sexual selection involving female mate choice. Traditionally, STIs were thought to select for monogamy (Immerman, 1986; Immerman and Mackey, 1997) but subsequent models showed that STIs can maintain both monogamy and promiscuity in a population, as well as select for risky female choice (Thrall et al., 1997; Boots and Knell, 2002; Kokko et al., 2002). (Female choosiness based on attractiveness of males is considered a risky strategy because the most popular males have the highest mating rate and are most likely to be infected.)

By altering the mating system, the STI changes its own ecological landscape, possibly setting the stage for coevolution. The outcome of coevolution can be hard to intuit. Indeed, there are several examples in the host-parasite literature of coevolution leading to different or unexpected outcomes compared to the evolution of either host or parasite in isolation (Gandon et al., 2002; Day and Burns, 2003; Best and White, 2009). Of particular interest is a model that considers the coevolution of host mate choosiness with virulence of an STI that affects host fecundity (Ashby and Boots, 2015). An investigation of STI virulence in the absence of host coevolution showed that a parasite that reduces the mating success of its host should evolve to be less virulent. Knell (1999) suggested that hosts would subsequently lose disease-avoidance behaviours such as mate choice based on the degree of parasitism of potential mates. However, when the level of host choosiness based on parasitism was allowed to coevolve with STI virulence, Ashby and Boots (2015) found that intermediate levels of disease-avoidance behaviour and virulence could evolve, and that coevolutionary cycling could occur between host choosiness and STI virulence. These unexpected results emphasize the importance of considering possible coevolutionary feedbacks of an STI with host mating system.

Over the last 25 years, it has become clear that, in many systems, sexual conflict over mating rate plays at least as large a role in shaping the evolution of male-female interactions as conventional sexual selection processes (Rice and Holland, 1997; Arnqvist and Rowe, 2005). In the absence of an STI, there are several possible outcomes of sexually antagonistic coevolution depending on the biology of the system. If male persistence and female resistance carry no inherent cost, traits will continually escalate in an evolutionary arms race (Gavrilets and Hayashi, 2006). Incorporating natural selection against persistence and resistance traits prevents runaway evolution (Gavrilets et al., 2001) and allowing for the evolution of female sensitivity can lead to female indifference to male traits, halting the coevolutionary process (Rowe et al., 2005). In all of these cases, only females suffer the cost of mating. Given that a sexually transmitted infection increases the cost of mating to both males and females it is not obvious how an STI will affect sexually antagonistic host interactions. Though classic theory indicates that the ESS virulence of an STI is unaffected by mating rates, there remains the potential for host-parasite coevolution. Because the traits mediating sexual conflict are themselves costly, evolutionary changes in these traits driven by the emergence of an STI might create epidemiological feedbacks that drive subsequent STI evolution.

We model an STI in a host system with sexual conflict over mating rate., i.e., the STI can coevolve with sexually antagonistic host traits (male persistence and female resistance). We examine how an STI changes the escalation of sexually antagonistic traits in the host, as well as how the evolution of these host traits affects whether an STI can establish and, if so, the evolution of STI virulence. The fitness of a host is determined by both its viability and its reproductive output whereas these two fitness components of the host are not equally important to the fitness of a sexually (horizontally) transmitted infection. For this reason, we consider separately cases where the fitness costs of the host’s sexually antagonistic traits occur through viability versus reproduction. In addition, we separately examine an STI that reduces host fitness via mortality and one that reduces sexual fitness (i.e., female fecundity and male mating success); we refer to these as “viability-reducing STI” and “reproduction-reducing STI” models, respectively. We highlight how the evolutionary outcomes that occur in cases where costs of sexually antagonistic traits and/or STI virulence manifest through host mortality contrast with cases where they manifest through host reproduction.

## Model Setup

We take as our focal case the model with a viability-reducing STI where the costs of sexually antagonistic (SA) traits are manifest through reductions in viability (hereafter “viability-reducing STI” and “viability costs for SA traits”). Other cases are subsequently described with respect to how they differ from this focal case. The results presented are from individual-based simulations. Numerical solutions of an analytical model of the focal case are similar (Supplementary Material).

### Host Life Cycle without the STI

We model an interlocus sexual conflict over mating rate by assuming sex-limited expression of male and female traits (each controlled by a single additive diploid locus). A male is characterized by his persistence trait and a female her resistance trait; these traits may be morphological (e.g., male grasping and female anti-grasping traits of water striders; Arnqvist and Rowe 2002) or behavioural (e.g., harassment by males or vigorous struggles by females in water striders; Arnqvist 1989). Resistance (*x*) and persistence (*y*) levels expressed by a host are calculated as the average of the trait values from each chromosome (e.g., *x* = (*x*_1_ + *x*_2_)/2 where *x*_1_ is the resistance allele on chromosome one and *x*_2_ is the resistance allele on chromosome two). Definitions of key parameters are provided in Table 1.

**Table 1.** Parameters and Variables

| Parameter | Definition |
| --- | --- |
| $y$ | male persistence trait |
| $x$ | female resistance trait |
| $u = y - x$ | mating rate metric |
| $\phi[u]$ | mating probability function |
| $c$ | persistence cost to males |
| $\delta$ | resistance cost to females |
| $d$ | mating cost to females |
| $\alpha$ | number of males encountered |
| $b$ | maximum fecundity |
| $K$ | carrying capacity |
| $M$ | number of males in the population |
| $F$ | number of females in the population |
| $m$ | number of males a given female mates with |
| $\mu_i$ | baseline mortality coefficient of sex $i$ |
| $v$ | virulence of the STI |
| $\beta[v]$ | STI transmission function |
| $w$ | transmission-virulence trade-off parameter |

Each breeding season, a female encounters a certain number of males, randomly drawn from a Poisson distribution with mean *α*. The probability of mating during an encounter between a male with persistence *y* and a female with resistance *x* is an increasing function of the difference *u* = *y* – *x* as given by *ϕ*[*u*] = 1/(1 + *e*^−*u*^) (following Gavrilets et al. 2001 and Rowe et al. 2005). The mating rate *ϕ*[*u*] plays several roles in the model. Importantly, it affects female mortality as females pay a viability cost *d* per mating. The mating rate also affects female fecundity in that a given female can have zero fitness if she remains unmated (at low mating rates) by the end of the breeding season. A mated female has maximum fecundity *b* (per breeding season) that is decreased by density-dependence if population size exceeds the carrying capacity *K*.

Mortality occurs after mating but prior to offspring production. Males and females suffer baseline mortality rates, *μ_m_* and *μ_f_*, respectively, and face nonlinearly increasing costs of expressing their respective sexually antagonistic traits. A male pays a cost for his persistence trait; the strength of this cost is *ce^y^* where c is the “persistence cost” parameter. Likewise, a female pays a mortality cost for her resistance trait given by *δe^y^* where *δ* is the “resistance cost” parameter. Viability selection against these sexually antagonistic traits, together with the cost of mating experienced by females, results in three costs incurred by hosts (persistence costs to males as well as resistance and mating costs to females). After mating (but before giving birth), adult females die with probability (1 – *e*^−(*μ_f_* + *δe^x^* + *dm*)^) and adult males die with probability (1 – *e*^−(*μ_m_* + *ce^y^*)^).

Surviving females produce offspring. The number of offspring born to a mated female is drawn from a Poisson distribution with mean *b*(1 – (*M* + *F*)/*K*) where *M* and *F* are the number of adult males and females in the population. If a female has mated with multiple males (let m be the number of males she mated with), a given male sires an average of 1/*m* of her offspring, with the actual sire of each offspring chosen at random from the m males. Gametes are formed with free recombination between loci and alleles experience mutation with probability *U*_hosts_ per locus. Effect size of a mutation is chosen from a normal distribution with a mean of zero and standard deviation of 0.1. If the mutational step yields a negative trait value, the trait value is assumed to be zero. The surviving adults and newborn offspring make up the next generation, whose mean and variance in host trait values are recorded before undergoing another round of selection, mating, and mutation.

### Inclusion of a Viability-Reducing STI

If an STI is present in the population, it can be transmitted from an infected individual to an uninfected (“susceptible”) individual with probability *β*[*v*], given a mating between them. *α*[*v*] is a function of STI virulence, v, and takes the standard form *β*[*v*] = *v*/(*w* + *v*), where *w* determines the shape of the trade-off between transmission and virulence (see Otto and Day, 2007, Chapter 12). An infected host suffers additional mortality *v*, e.g., adult males die with probability (1 – *e*^−(*μ_m_*+*ce^y^*+*v*)^). A newly infected host cannot infect other hosts during the same breeding period in which it became infected itself, i.e., the newly infected host must survive to the next breeding period before it is infectious. During this latent period, the STI undergoes mutation with probability *U*_STI_. Effect size of a mutation is chosen from a normal distribution with a mean of zero and standard deviation of 0.01. If the mutational step yields negative virulence, virulence is assumed to be zero. At the start of each generation, the population mean and variance in virulence are recorded.

### Reproduction Costs for SA Traits

Simulations proceed as above except the cost of sexually antagonistic traits affects aspects of reproduction instead of viability. Higher resistance trait values reduce female fecundity such that the number of offspring born to a mated female is drawn from a Poisson distribution with mean *e*^−*δe^x^*^ *b*(1 – (*M* + *F*)/*K*). Higher persistence trait values reduce male fertility by decreasing siring success (i.e., the male’s persistence trait increases his expected number of matings but decreases his postcopulatory competitive ability). Specifically, having mated with a given female, a male sires an average of 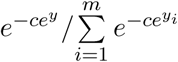 of her offspring, where *m* is the total number of males she mated with and *y_i_* is the persistence trait of male *i*.

### Reproduction-Reducing STI

Simulations with a reproduction-reducing STI proceed as described for a viability-reducing STI except disease virulence affects host fitness through aspects of reproduction. Female fecundity is reduced by infection such that the number of offspring born to a mated female is drawn from a Poisson distribution with mean *e*^−*v*^ *b*(1 – (*M* + *F*)/*K*). To prevent continually escalating virulence evolution, a reproduction-reducing STI faces a transmission-virulence trade-off in males. Infected males have a lower probability of mating with an encountered female than an uninfected male with the same persistence trait value; infected males can be considered as exhibiting lower mating effort than uninfected males. Specifically, the probability of mating with an infected male is *e*^−*v*^ *ϕ*[*u*]. Note that like a viability-reducing STI, there is a latent period before an individual is infectious.

### Initial Conditions

Individual-based simulations were carried out in Python (available upon request) for 50,000 generations, well after the trait values appeared to reach evolutionary equilibrium. To initiate the population, diploid hosts are assigned trait values at the resistance and persistence loci by drawing random values from a normal distribution with mean 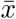 and 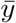, respectively, and standard deviation 0.5. In host populations infected with a sexually transmitted infection (STI), the STI is introduced into 5% of hosts with the virulence of each infection drawn from a normal distribution with mean 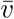 and standard deviation 0.1. Values from the last 1000 generations were averaged and reported in Figs. 1 and 4. Rare stochastic extinctions of the host population are excluded from the average.

**Figure 1:**
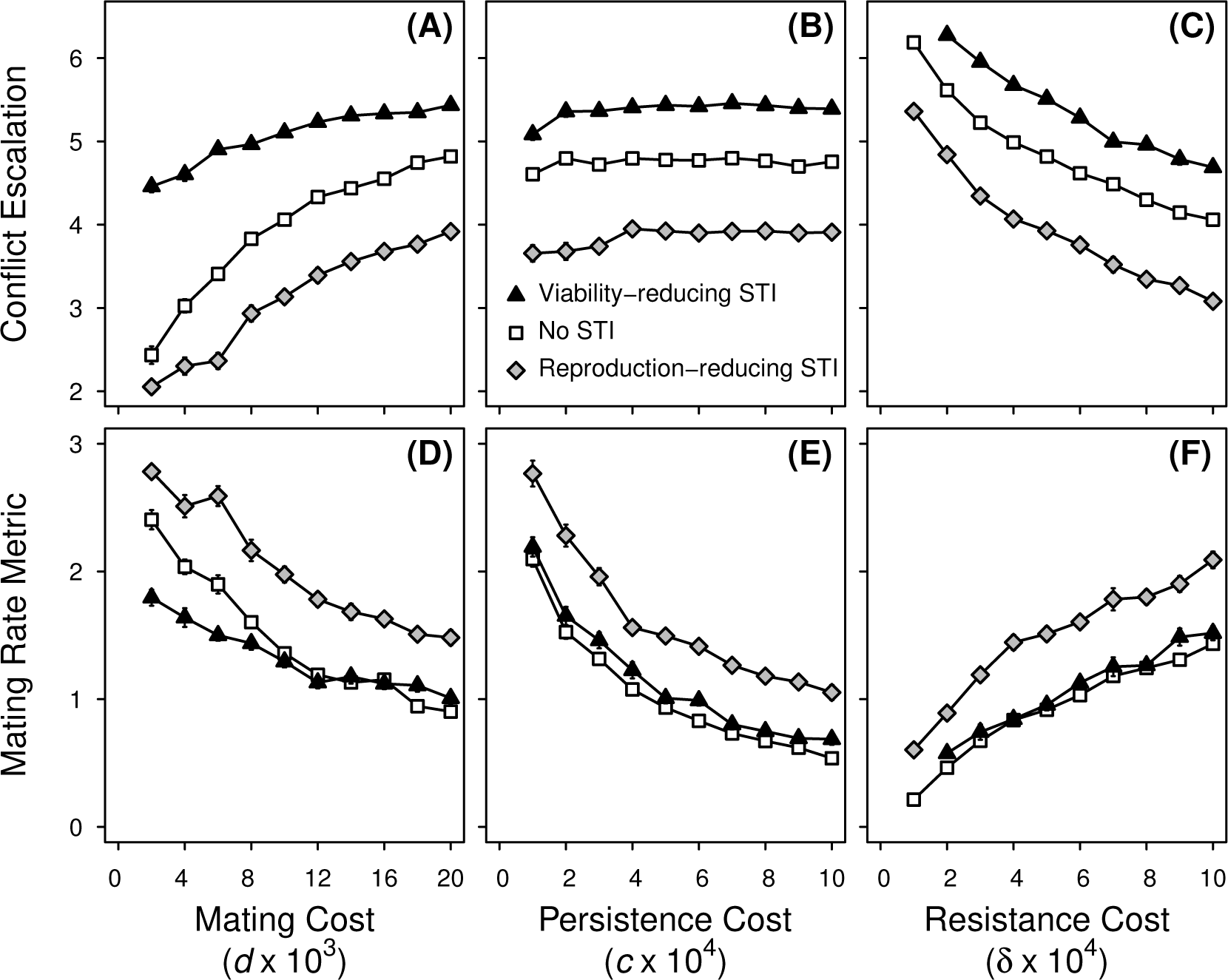
The outcome of coevolution between males and females in the absence of an STI and in the presence of two types of evolving STIs. Both sexes pay viability costs for expressing sexually antagonistic traits. Shown here are the degree of escalation in sexual conflict, measured as the average of *y* and *x* (*A* - *C*), and the difference between persistence, *y*, and resistance, *x* (*D* - *F*); values are at evolutionary equilibrium. The mating rate is an increasing function of the difference between male persistence and female resistance (*y* – *x*); thus, we label this difference the ‘mating rate metric’. Simulations with an evolving STI began at the STI-absent host trait values and the results for a reproduction-reducing STI (diamonds) and a viability-reducing STI (triangles) are a three-way evolutionary equilibrium with the STI at evolutionary equilibrium virulence *v* (see Fig. 4A). Parameter values (unless shown otherwise): *α* = 10, *μ* = 0.2, *b* = 4, *K* = 1000, *w* = 1, with *c* = 0.0005, *δ* = 0.0005, *d* = 0.02. Each point represents the average of 20 independent simulations +/- standard error. Simulations in which the disease went extinct are excluded from calculating the average. At very low resistance costs, a viability-reducing STI went extinct in 100% of simulations (see Fig. 3) and there is no data for the three-way evolutionary equilibrium. Qualitatively similar patterns are observed when hosts pay reproduction costs for expressing sexually antagonistic traits (see Supplementary Material Fig. S1).

## Results

### Evolution of Host Traits without an STI

To understand how a coevolving STI affects the outcome of sexually antagonistic coevolution we first consider how the host evolves in the absence of an STI (which has been previously studied for ‘mortality costs’ of sexually antagonistic (SA) host traits, e.g., Gavrilets et al. 2001). Rather than reporting the SA traits directly, it is more useful to consider two other metrics instead. The average of the two trait values, (*y* + *x*)/2, is an indicator of the degree of escalation in the SA traits (hereafter, “conflict escalation”) and is plotted in Fig. 1A-C. Second, we plot the difference between male persistence and female resistance, *u* = *y* – *x* (hereafter, “mating rate metric”) because the mating rate is an increasing function of this difference (Fig. 1D-F). The evolutionary equilibrium trait values presented in Fig. 1 and all subsequent figures are stable to different initial values.

For the range of parameters shown in Fig. 1, male persistence *y* always exceeds female resistance *x*, so the mating rate metric *u* is positive and mating rates are not too low. The extent of conflict escalation, as reflected by average trait values, increases substantially as the mating cost to females increases (Fig. 1A). This is driven by the female trait increasing to reduce the mating rate, while the male trait increases only slightly in response because of the nonlinearly increasing cost of natural selection. Because female resistance increases more than male persistence, the mating rate metric *u* declines (Fig. 1D) with increased mating cost. Conflict escalation does not change drastically with increasing cost of male persistence (Fig. 1B) but there is a reduction in the mating rate (Fig. 1E). In contrast, increasing the cost of female resistance leads to a decline in conflict escalation (Fig. 1C) accompanied by an increase in the mating rate metric (Fig. 1F). At high resistance costs, females with lower resistance better balance the costs of mating with the costs of their SA trait. Qualitatively similar patterns are observed when the costs for sexually antagonistic traits occur through reproduction (Fig. S1) rather than viability (Fig. 1), although there is a noticeable decrease in the mating rate metric.

### Evolution of Host Traits with an Evolving STI

#### Viability-Reducing STI

We now consider the evolution of host traits in the presence of an evolving sexually transmitted infection and then later discuss virulence of the STI. We begin by examining a viability-reducing STI in a host where the costs of SA traits occur through viability effects. The introduction of an evolving viability-reducing STI escalates the conflict between the sexes (Fig. 1). (A viability-reducing STI also causes an increase in conflict escalation if costs of SA traits occur through reproduction, though the increase is smaller; Fig. S1.) This increase is driven by females. If only the females are allowed to evolve (not shown), female resistance will evolve to be higher than male persistence to reduce the additional cost of mating arising from the risk of STI infection. In this case, *u* = *y* – *x* is negative and the mating rate drops to low levels. A viability-reducing STI cannot persist at low mating mates. In contrast, if only the males are allowed to evolve (not shown), male persistence increases from its STI-absent equilibrium, presumably because males are selected to more quickly obtain additional mates in the face of higher mortality rates due to infection (though the change is smaller compared to the case where females evolve). The STI does not go extinct because mating rates remain high (as *u* increases). When females and males can both evolve, the increase in female resistance invokes a subsequent increase in male persistence so males ensure mating (Fig. 2A). As such, the net effect of coevolutionary feedback tends to be increased average trait values (i.e., increased conflict escalation). It is worth emphasizing that the evolution of male persistence in response to the escalation in female resistance allows the STI to remain in the system. If a lack of genetic variation in the male trait prevented its coevolution, the STI would go extinct after female resistance increased and caused a decrease in mating rates.

**Figure 2:**
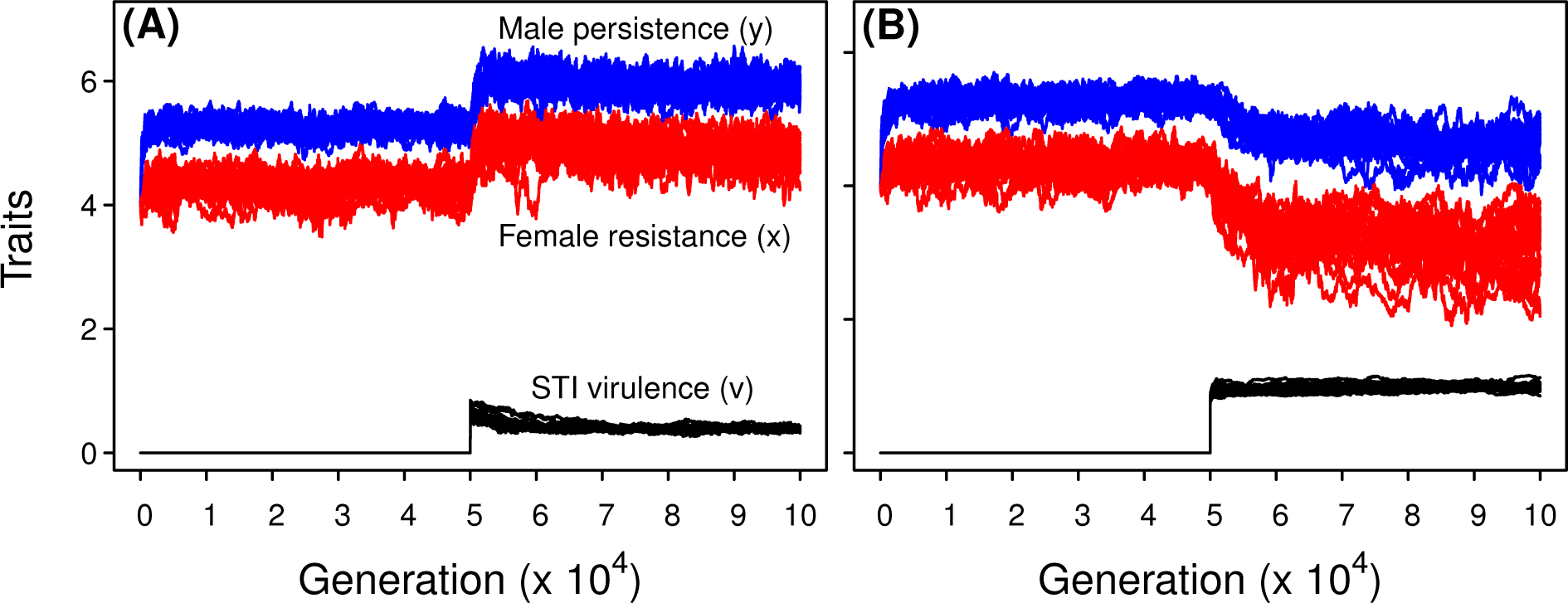
Sample runs from individual-based simulations (20 replicates) where hosts evolve to their disease-absent equilibrium values before a viability-reducing STI (A) or a reproduction-reducing STI (B) is introduced. Male persistence is shown in blue, female resistance in red, and STI virulence in black. Parameter values: *α* = 10, *μ* = 0.2, *b* = 4, *K* = 1000, *w* = 1, *d* = 0.02, *c* = 0.0005, *δ* = 0.0005.

#### Reproduction-Reducing STI

A reproduction-reducing STI has qualitatively different consequences for the outcome of sexual conflict. Again, this outcome can be understood by first thinking about the evolution of each sex in isolation. If only females are allowed to evolve (not shown), female resistance decreases in the presence of a reproduction-reducing STI. A prevalent STI results in less mating pressure on females because infected males exhibit reduced mating effort compared to uninfected males. Consequently, females do not need to invest as much in costly resistance traits. If only males are allowed to evolve, there is no observed change in the male persistence trait. When males and females can both evolve, the decrease in female resistance allows male persistence to decrease slightly (although to a lesser extent). Thus, in contrast to a viability-reducing STI, a reproduction-reducing STI decreases the degree of conflict escalation (Fig. 1A-C, Fig. 2B). This is also true if the costs of the SA traits occur through reproduction (Fig. S1) rather than viability (Fig. 1). In contrast to a viability-reducing STI, evolution of either host sex never drives a reproduction-reducing STI extinct for the parameters examined here.

#### Establishment of the Sexually Transmitted Infection

In the section above, we focused on cases where both hosts and parasites can evolve and the disease establishes itself within the host population. (We use the term disease “establishment” rather than the more typical term “persistence” to avoid confusion with the host male’s “persistence” trait.)

We now focus on understanding how a lack of genetic variation that prevents evolution of either host or parasite traits affects both the establishment and virulence of the STI. We discussed one important case above in which a lack of genetic variation in the male trait but not the female trait prevents the establishment of an evolving viability-reducing STI. Here we focus on contrasts between hosts and parasites in their evolvability (rather than on differences in evolvability between male and female host traits).

When neither the host nor parasite can evolve, a viability-reducing STI near its equilibrium virulence always establishes when introduced into host populations at equilibrium except for rare stochastic extinctions (for the parameter values considered here). In comparison, a relatively avirulent viability-reducing STI almost always goes extinct and a highly virulent one (shown in Fig. 3, open symbols) establishes unless mating rates are very low (i.e., when mating costs are high and there are high persistence costs to males or low resistance costs to females). In contrast, in all cases examined, a reproduction-reducing STI establishes regardless of whether it was introduced at low, high, or equilibrium virulence. By virtue of not killing its host, a fertility STI has a long duration of infection and can establish and spread at very low mating rates through rare transmission events. Because of the ease of establishment of a reproduction-reducing STI (for all combinations of evolvability), further discussion of extinction rates and the accompanying figure (Fig. 3) pertain only to a viability-reducing STI.

**Figure 3:**
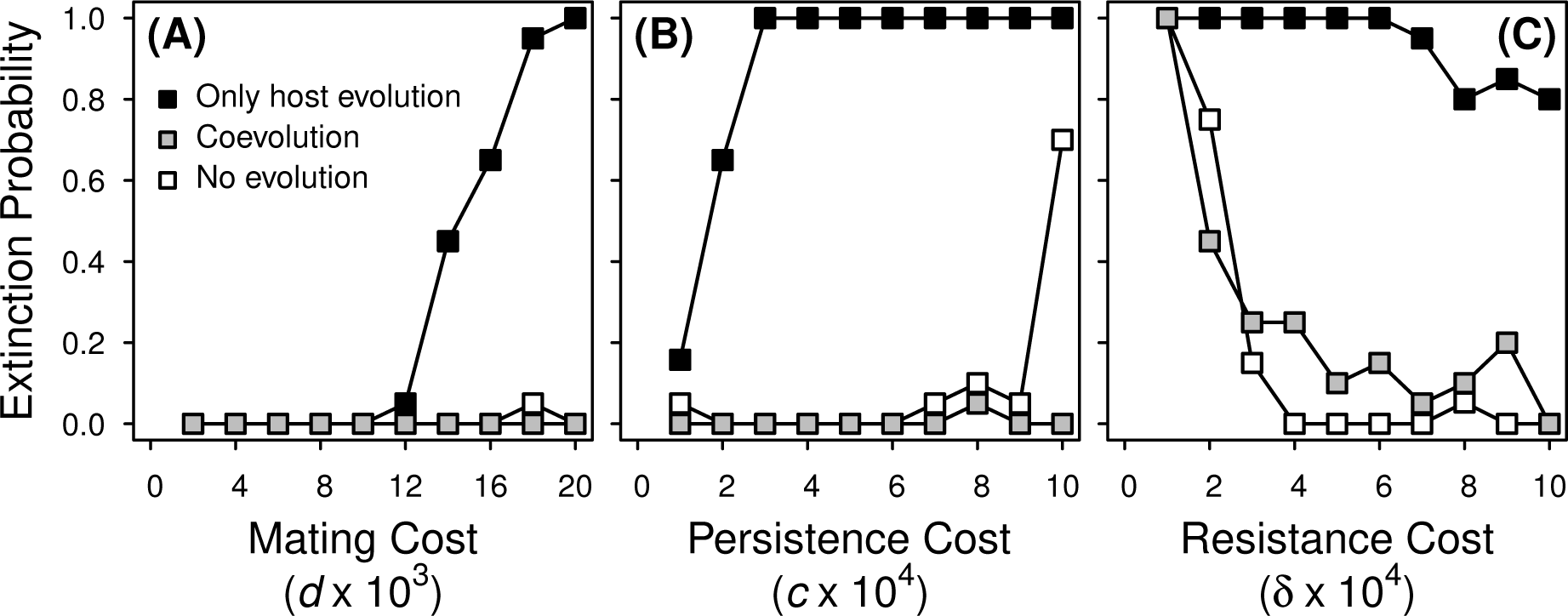
Fraction of simulation runs where a highly virulent viability-reducing STI was driven extinct in hosts paying viability costs. The parasite was introduced with *v* = 0.8 at the STI-absent male persistence and female resistance host trait values and neither host nor parasite evolved (open symbols), host traits evolved in the presence of a non-evolving parasite (black symbols), or hosts and STI both evolved (gray symbols). Parameter values (unless shown otherwise): *α* = 10, *μ* = 0.2, *b* = 4, *K* = 1000, *w* = 1, with *c* = 0.0005, *δ* = 0.0005, and d = 0.02. The extinction probability was determined from 20 independent simulations. STI extinction probability was slightly higher, on average, in hosts paying reproduction costs for sexually antagonistic traits, but overall patterns of extinction probability were qualitatively similar (Supplementary Material Fig. S1).

If the host but not the parasite can evolve, then females are selected to increase their SA resistance trait in the presence of a viability-reducing STI, driving an increase in average host trait values and a decrease in the difference between the sexes (not shown). The accompanying decrease in mating rate can drive a highly virulent, non-evolving STI extinct (i.e., at high persistence costs to males, low resistance costs to females, and high mating costs to females). In Fig. 3 we show the fraction of runs in which the parasite went extinct when hosts can evolve but parasites cannot (black symbols). Extinction rates are higher over a wider range of parameter values when female resistance costs are low (*δ* = 0.0001, data not shown). As noted above, STI extinction is even more likely if the female trait can evolve but the male trait cannot. For any of the parameter values shown in Fig. 3 a viability-reducing STI is unable to invade if only the female host trait evolves (not shown). If the STI can evolve but hosts cannot evolve, an initially virulent viability-reducing STI only goes extinct for very low mating rates (not shown).

Finally, we consider evolution in both the host and the STI (gray symbols, Fig. 3). STI extinction occurs most often when the cost of resistance to females is low (Fig. 3C). At low resistance costs, elevated female trait values quickly evolve in the presence of the STI, which lowers the mating rate enough (albeit, transiently) that a viability-reducing STI goes extinct due to lack of transmission opportunity. When the STI can coevolve with an evolving host, it is likely to successfully establish across a wide range of conditions under which it would otherwise go extinct (Fig. 3). Contrasting the case where the costs of sexual antagonism are paid through viability (Fig. 3) rather than reproduction (Fig. S2), the STI is more likely to go extinct in the latter scenario for most parameter values. Differences between these scenarios in STI establishment probability are likely driven by the pronounced decrease in mating rate accompanying the introduction of the STI in host populations paying costs for SA traits through reproduction (compare Fig. 1D-F to Fig. S1D-F).

#### Virulence of a Viability-Reducing STI

Conditional on establishment of a viability-reducing STI, three-way evolution of the STI with male and female host traits can give rise to quantitatively different results than if the STI was introduced into a non-evolving host population (Fig. 4A-C). A viability-reducing STI becomes more virulent if hosts coevolve than if they do not, but only if the cost of SA traits occur through viability. This is because the addition of the STI to the system causes evolutionary increases in female resistance and, consequently, male persistence, thereby increasing the host mortality rate; optimal STI virulence is expected to increase with host mortality rate (Knell, 1999). The equilibrium virulence of a viability-reducing STI increases as the average host trait values increase and does not depend on the difference in trait values. The difference in trait values only affects the establishment and prevalence of the STI and not its evolutionary equilibrium virulence. In contrast, when the cost of SA traits occur through reproduction, the virulence of a viability-reducing STI does not increase with an evolving host (Fig. 4A-C) even where there is host conflict escalation in the presence of the STI (Fig. S1A-C). Escalated male and female trait values do not increase host mortality and invoke subsequent coevolution of the STI.

**Figure 4:**
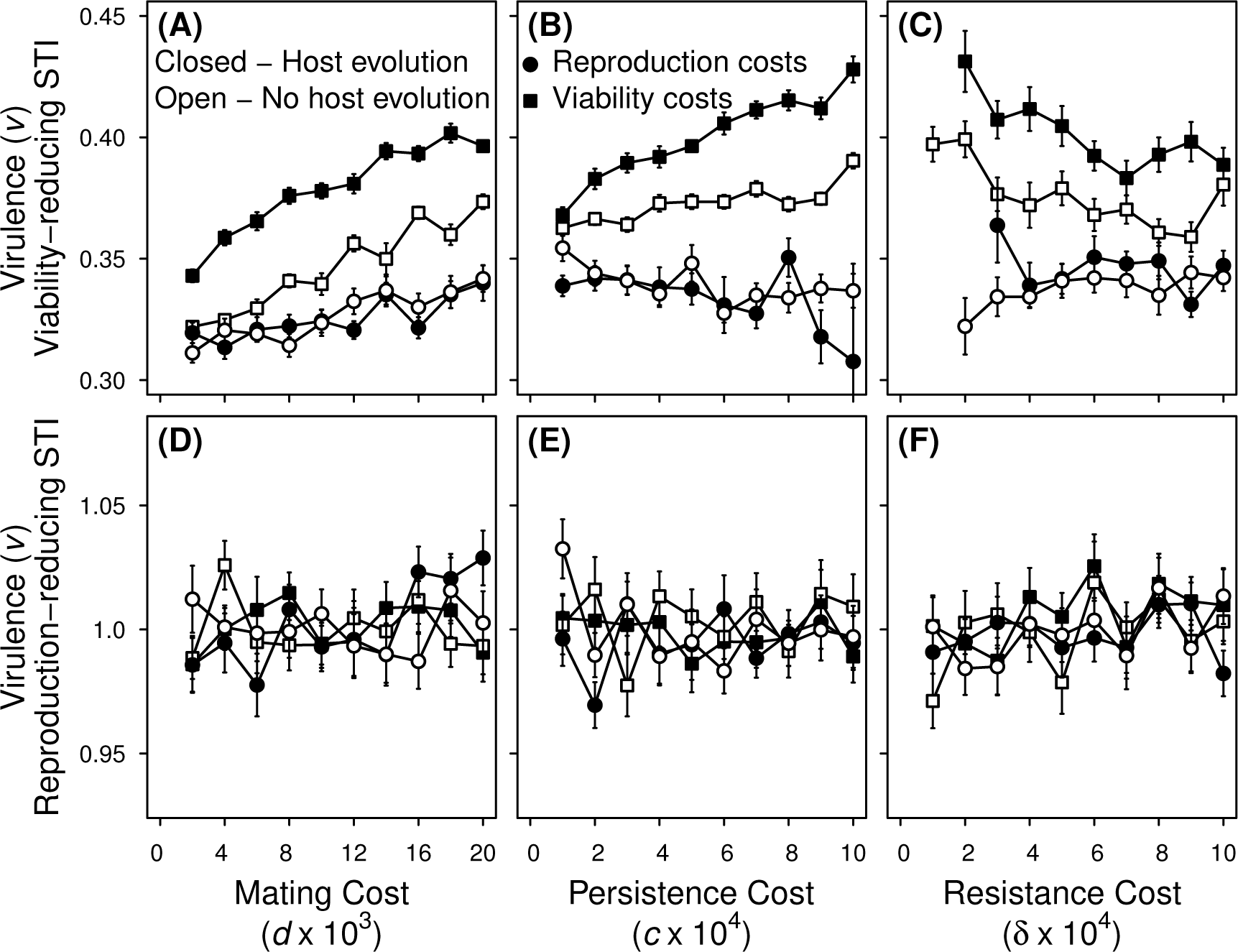
Evolutionary equilibrium virulence of two types of evolving STIs in a population where hosts do not evolve (open symbols) or do evolve (closed symbols). Virulence of a viability-reducing STI (A-C) and a reproduction-reducing STI (D-F). In both cases, the parasite was introduced with *v* = 0.8 at the STI-absent male persistence and female resistance host trait values. Parameter values (unless shown otherwise): *α* = 10, *μ* = 0.2, *b* = 4, *K* = 1000, *w* = 1, with *c* = 0.0005, *δ* = 0.0005, *d* = 0.02. Each point represents the average of 20 independent simulations +/- standard error. Simulations in which the disease went extinct are excluded from calculating the average and can lead to larger error bars at high extinction probabilities. At very low resistance costs a viability-reducing STI infecting hosts paying viability costs went extinct in 100% of simulations (see Fig. 3) and there is no data for the three-way evolutionary equilibrium.

#### Virulence of a Reproduction-Reducing STI

Virulence evolution of a reproduction-reducing STI is shaped by its effects on males. In infected females, a reproduction-reducing STI does not face a transmission-virulence trade-off because the consequences of infection (decreased fecundity) do not reduce transmission opportunities. However, infected males in our model suffer reduced mating success with females. Thus, males infected with a highly virulent reproduction-reducing STI do not mate as often as uninfected males with similar persistence trait values. This reduction in transmission opportunities creates a transmission virulence trade-off in one sex (males) that limits virulence evolution. The transmission-virulence trade-off is not affected by the host trait values themselves, regardless of whether costs of SA traits are paid through viability or reproduction. Consequently, the evolution of sexually antagonistic host traits induced by the presence of a reproduction-reproducing STI does not invoke a corresponding response in STI virulence evolution. In the absence of a coevolutionary response, virulence of a reproduction-reducing STI is the same in host populations that can and cannot evolve (Fig. 4D-F).

#### Comparison with a Horizontally Transmitted Disease

In our focal case, a viability-reducing STI coevolves with a host population paying SA costs through viability. As noted, STI virulence evolution depends on host mortality much like a horizontally transmitted disease or “ordinary infectious disease” (OID). Furthermore, STI virulence in our model causes disease-induced mortality. Given that both STI and OID virulence affect viability and are shaped by the same parameters, we might expect similar co-evolutionary outcomes if we model an OID infecting hosts with sexual conflict over mating rate. An OID, however, is not transmitted sexually and therefore will not exert direct selection on females to reduce the mating rate. We find that an OID has the same effect on the outcome of the conflict as increasing the baseline mortality rate *μ* (not shown). In contrast to a viability-reducing STI, an OID has little effect on the extent of conflict escalation. However, an OID drives an increase in the mating rate metric because male persistence increases and female resistance decreases.

## Discussion

Although the evolution of STI virulence and sexual conflict have each been studied in isolation (e.g., STIs: Knell 1999; Lipsitch and Nowak 1995, sexual conflict: Gavrilets et al. 2001; Gavrilets and Hayashi 2006), the link between them has received little attention. The models presented here are aimed at understanding this connection. In some cases, each species reciprocally affects the evolution of the other (i.e., true coevolution) whereas in other cases only one species evolves in response to the other. Whether, and in which direction, each species evolves in response to the other depends on how trade-offs are structured.

We found that a sexually transmitted infection affects the level of escalation of traits mediating sexual conflict within the host, but that viability-reducing STIs and reproduction-reducing STIs do so in opposite ways. The introduction of a viability-reducing STI escalates the conflict. Female resistance increases to reduce the additional cost of mating imposed by the risk of disease infection. Male persistence increases to stay above increasing female resistance levels. Thus, average host trait values increase, i.e., there is elevated conflict escalation. If the cost of SA traits occurs through viability, then a co-evolving viability-reducing STI increases its virulence level in response to these changes in the host, not because of a change in host mating rate but rather because of the increased host death rate resulting from the escalated sexual conflict. In contrast, if the cost of SA traits occurs through reproduction, then increased conflict escalation evolves in response to a viability-reducing STI but in this case there is no feedback affecting virulence evolution.

In both cases described above, the three traits reach a stable equilibrium. We find no evidence of cycling as has been observed under some conditions in a coevolution STI model involving conventional sexual selection Ashby and Boots (2015). Based on our understanding of why the observed changes occur, cycling is not expected. Stabilizing selection maintains the evolutionary equilibrium. Male persistence increases male reproductive success at the expense of other costs associated with persistence such as increased predation risk (Rowe, 1994) or reduced foraging time (Robinson and Doyle, 1985). Females suffer costs of mating but must balance these with the direct cost of expressing the resistance trait and the risk of remaining unfertilized if resistance is too high. In the presence of a viability-reducing STI, males experience stronger selection to obtain mates quickly in the face of higher total mortality and females experience stronger selection to reduce the additional cost of mating associated with a prevalent STI. The escalation of sexual conflict increases total host mortality, selecting for higher STI virulence, which in turn should drive further escalation of the sexual conflict traits. However, the “faster than linear” increasing costs to hosts of the escalating sexual conflict traits and the “slower than linear” increasing transmission benefits of increased STI virulence ensures that the system reaches an equilibrium rather than evolving to ever higher levels of all three traits. (Such non-linear costs are a common assumption in both sexual conflict and virulence models and are necessary to have sensible equilibria (e.g., Gavrilets et al. 2001 and Otto and Day 2007).) Evolutionary equilibrium is even easier to understand when hosts pay costs of SA traits through reproduction because then conflict escalation does not select for higher STI virulence and create the potential for a host-STI arms race.

Compared to viability-reducing STIs, reproduction-reducing STIs have the opposite effect on the outcome of sexual conflict: a reproduction-reducing STI de-escalates the conflict. Females evolve lower resistance in response to the alleviation in mating pressure they experience because infected males have reduced mating success. The accompanying (but smaller) decline in the male trait contributes to conflict de-escalation. Regardless of whether hosts pay the cost of SA traits through viability or reproduction, virulence of a reproduction-reducing STI does not seem to be affected by the decrease in average host trait values. All three traits reach evolutionary equilibrium due to stabilizing selection. Continual de-escalation of the conflict is prevented by selection on females to resist costly matings and selection on males to stay competitive given they will enter fewer matings if they are infected.

When the cost of SA traits occur through viability, conflict escalation occurs in response to either a viability-reducing STI or a reproduction-reducing STI. However, a coevolutionary response from the STI occurs only with a viability-reducing STI. As known from past virulence theory (Anderson and May, 1982; Ewald, 1983), the transmission-virulence trade-off can be thought of as a trade-off between current and future transmission for the STI, the latter of which requires the current host’s survival. In cases where host mortality is higher for reasons not directly due to the disease (e.g., higher extrinsic mortality or higher investment into SA traits carrying viability costs), then future transmission is down weighted and the disease evolves higher virulence (i.e., increased investment in current transmission). This logic does not apply to a reproduction-reducing STI. As modeled here, the transmission-virulence trade-off in a reproduction-reducing STI occurs because increased transmission *given a mating* comes at the expense of reducing a male’s probability of successfully obtaining a mate. That is, the two components of this trade-off both affect current transmission. Thus, their value relative to one another is unaffected by extrinsic factors that alter the value of future transmission.

One of the costs we did not explore in depth is the risk of females remaining unfertilized. In the simulation model, a female achieved her full fecundity provided that she mated with at least one male. However, if fertilization was not guaranteed by mating, the increased risk of being left unfertilized would be expected to affect the evolution of female resistance. Thrall et al. (1997) constructed a model investigating how male and female mating behaviour affected reproductive success in the presence of an STI. At high disease prevalence, females could achieve the same fitness by being monogamous and minimizing infection risk or by being promiscuous and minimizing the risk of being left unfertilized but increasing the risk of infection. It is possible that if there were higher probabilities of females being unfertilized in our model, divergent female strategies (low and high resistance) would be maintained.

We investigated the full three-way evolutionary interaction over a range of mating costs to females. However, many empirical investigations of sexual conflict in natural systems have reported females suffer a cost of harassment instead of, or in addition to, the cost of mating (Alcock et al., 1977; Rowe, 1994; Stone, 1995; Jormalainen, 1998; Watson et al., 1998). At high male densities or male-biased sex ratios, the cost of rejecting harassing males can become so great that females are selected to decrease resistance, increasing overall mating activity in these systems (Rowe, 1992; Rowe et al., 1994; Lauer et al., 1996). However, the introduction of a STI would effectively increase the cost of mating, possibly tipping the balance in favour of high resistance for females.

The majority of models that have investigated the evolution of host mating strategies in the presence of an STI have assumed there is sexual selection, but no sexual conflict. The distinction between male attractiveness and female preference versus male persistence and female resistance has important consequences for the evolution of an STI. Choosiness may help females gain indirect benefits from mating with a preferred male. Resistance on the other hand helps females avoid the direct costs of mating (Gavrilets et al., 2001). Unless indirect benefits are strong (Thrall et al., 1997; Boots and Knell, 2002), the presence of an STI will cause selection for non-choosy females because the most popular males have high infection prevalence (Thrall et al., 2000). In a system with sexual conflict, those males with the highest persistence traits would be more likely than average males to be infected, adding to the cost of mating for females. Females do not reduce their risk of mating with such males by having lower resistance (i.e., lower resistance is not equivalent to being less choosy because the latter can mean less discriminant mating without changing mating frequency). A female with lower resistance may have a lower fraction of her matings with high persistence (probable STI-carrying) males but she will not have a fewer number of matings with such males. Moreover, she will have more total matings, increasing her infection risk. Consistent with this, we found that the presence of a viability-reducing STI resulted in increased female resistance, which might be construed as increased choosiness, opposite to what is observed in sexual selection models. On the other hand, a reproduction-reducing STI results in decreased female resistance, which might be construed as decreased choosiness. Though this would appear to match the outcome of sexual selection models, the reason for this result is very different. In the sexual conflict model, reduced female resistance evolves not to avoid infection but rather because the STI reduces the mating effort exhibited by infected males so females do not need to invest as heavily in their resistance trait.

For the most part, we have assumed that infection does not directly alter a male’s persistence or a female’s resistance. However, in some cases STIs manipulate host behaviour in ways that directly affect sexual conflict. An STI that influences male competitive ability could change the relative costs of mating. For example, parasitized males can be less competitive (Siva Jothy and Plaistow, 1999; Thomas et al., 1999), decreasing the risks of mating for females in a population with a highly prevalent STI. This is the scenario we modeled by allowing a reproduction-reducing STI to decrease mating probability for infected males. Conversely, as in the milkweed leaf beetle (Abbot and Dill, 2001), STI-infected males can be more aggressive than their uninfected counterparts, which could increase the number of matings and therefore the costs to females. Alternatively, the STI may manipulate host behaviour in such a way that its interests are aligned with one sex. It has been suggested that an STI that reduced female remating rate would be beneficial for males (Knell and Webberley, 2004) because a male that infects his mate would reduce her remating rate and ensure his own paternity. If the benefits of reducing sperm competition outweigh the costs of the STI, males may be selected to increase persistence and consequently their likelihood of acquiring an STI. Similarly, there is some evidence that infection by an STI increases oviposition rates in the fall army worm moth, meaning that males could benefit from acquiring and transmitting an STI (Simmons and Rogers, 1994). There are numerous ways males and females could evolve in response to these changes in host mating behaviour. Overall, these examples suggest that a change in the cost structure in the presence of an STI could affect the outcome of sexual conflict in natural systems.

There are many well-known examples of STIs and sexual conflict but we are aware of no systems i where both are well studied. Evidence of sexual conflict and sexually transmitted diseases has been reported in ungulates (conflict: Bro-Jørgensen 2010; STI: Lockhart et al. 1996), Drosophila (conflict: e.g., Rice et al. 2006; STI: Knell and Webberley 2004), and the two-spot ladybird *Adalia bipunctata* (conflict: Haddrill et al. 2013; STI: Webberley et al. 2002; Ryder et al. 2005). Studies of sexually antagonistic traits in populations infected with a viability- or reproduction-reducing STI should look for escalated or de-escalated trait values, respectively, compared to populations where the STI is absent. Research on viability-reducing STIs should compare different populations or closely related host species that experience different persistence, resistance, or mating costs. ‘ Variation in the costs of sexually antagonistic host traits could arise between populations if, for example, the persistence or resistance trait made the bearer more vulnerable to predation in an open versus closed habitat (see Fricke et al., 2009, for a discussion of the dependence of sexually antagonistic selection on environmental conditions). Higher mortality rates in one habitat should select for higher virulence in an endemic viability-reducing STI. Additionally, we expect lower disease prevalence in populations where male persistence is strongly selected against and there are high costs of mating or low costs of resistance for females, i.e., low mating rate will limit STI transmission.

We have shown that an STI has the potential to influence the outcome of sexually antagonistic coevolution. Because STIs are ubiquitous in nature (Lockhart et al., 1996), they should co-occur with sexual conflict often enough that it is worth considering how STIs change sexually antagonistic selection. Furthermore, considering the full coevolutionary interaction has important implications for both the susceptibility of a host population to invasion of a new STI and the level of virulence expected to evolve.

## Acknowledgements

This work was supported by the Natural Sciences and Engineering Research Council of Canada (Discovery Grant to AFA and Alexander Graham Bell Canada Graduate Scholarship to AMW). We would like to thank Samuel Alizon and Philip Greenspoon for comments on an earlier version of the manuscript.

## Supplemental Information

### Reproduction Costs for Sexually Antagonistic Traits

#### Analytical Model

We use a system of differential equations to describe sexually antagonistic coevolution in a population of haploid hosts (parameter definitions summarized in Table 1). Most parameters are the same as the individual-based simulations with the exception of *α*, which captures the sexual encounter rate as a fraction of the density-dependent interaction of males and females (instead of *α* being a fixed number of males that a female encounters in the computer simulation model). The probability of a female not mating is described by *γ*[*M*] = (1 – (*ϕ*[*u*])^*α*^*M*, where *ϕ*[*u*] = *y* – *x* (as above) and M is the number of males in the host population.

The ecological dynamics are captured by the following system of equations:

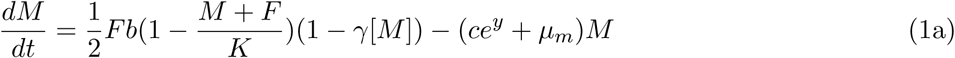

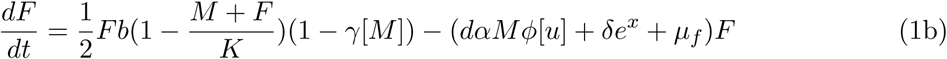

We find evolutionary stable (ESS) persistence and resistance, *y*\* and *x*\*, respectively, using sequential evolutionary invasion analyses for multiple traits. We plot the degree of conflict escalation, (*y*\* + *x*\*)/2, and the mating rate metric, *u*\* = *y*\* – *x*\*, for varying costs in Fig. S3.

We introduce a sexually transmitted infection into the model by dividing the host population into susceptible and infected classes. Susceptible individuals may contract the disease by mating with infected individuals of the opposite sex. We model transmission of the STI as density-dependent because it is reasonable to assume that the number of matings per capita will increase with density in systems governed by sexual conflict (i.e., at high density, females encounter more males and thus are likely to end up mating more total times holding *x* and *y* constant). To determine a female’s full cost of mating, we must track the expected number of males a female mates with and not just the fraction of her mates that were infected (as would be done under frequency-dependent transmission). Density-dependent sexual disease transmission has been documented in nature (for example in two-spot ladybird beetles, *Adalia bipunctata*, Ryder et al. 2005).

STI virulence, *v*, results in higher mortality of disease carriers. We assume a trade-off between transmission and virulence such that the transmission rate during mating is a saturating function of virulence *β*[*v*] = *v*/(*w*+*v*), where w determines the shape of the function (i.e., low values of w cause the transmission benefits of increased virulence to saturate more quickly). The epidemiological dynamics are described by the following set of differential equations:

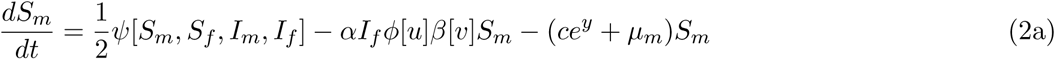

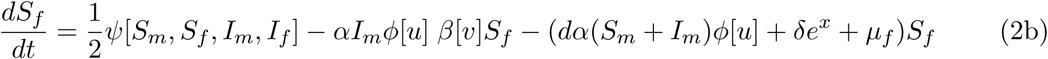

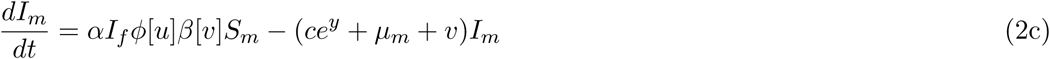

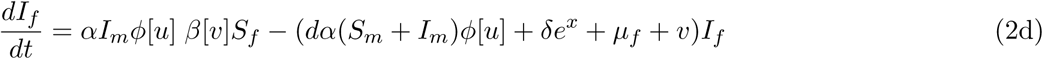

where

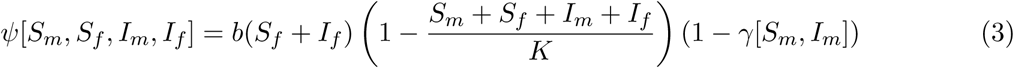

Similar to the STI-absent model, we find ESS persistence (*y*\*), resistance (*x*\*), and virulence (v*) using sequential evolutionary invasion analyses for multiple traits. We compare the degree of host conflict escalation and the mating rate metric to their respective values in the absence of an STI (Fig. S3). Finally, we compare STI virulence in a non-evolving host population (at its STI-absent ESS) to virulence in a coevolving host population (Fig. S4). Results are qualitatively similar to those of the individual-based computer simulations.

**Figure S1:**
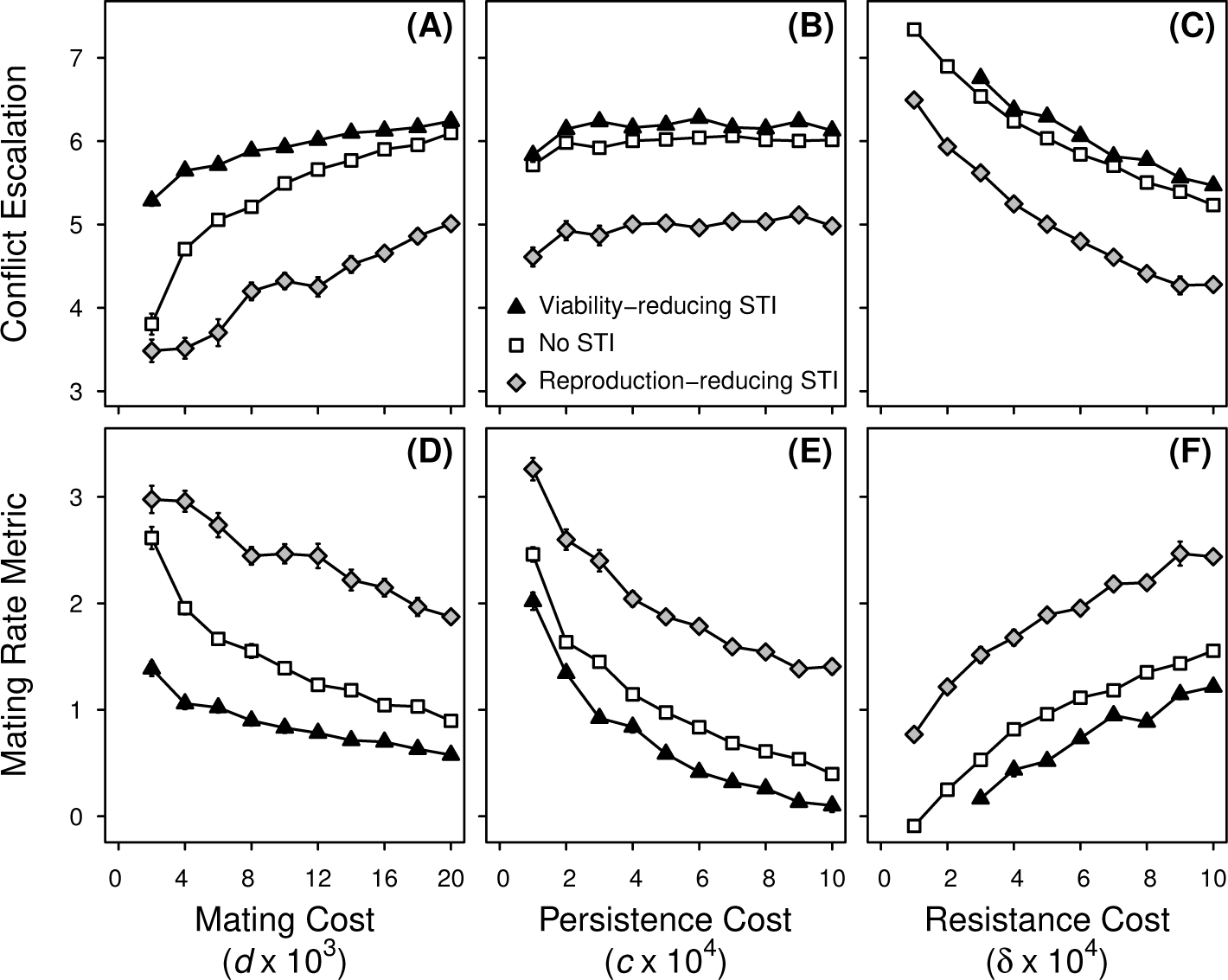
The outcome of coevolution between males and females in the absence of an STI and in the presence of two types of evolving STIs. Both sexes pay reproduction costs for expressing sexually antagonistic traits. Shown here are the degree of escalation in sexual conflict, measured as the average of *y* and *x* (*A* - *C*), and the difference between persistence, *y*, and resistance, *x* (*D* - *F*); values are at evolutionary equilibrium. The mating rate is an increasing function of the difference between male persistence and female resistance (*y* – *x*); thus, we label this difference the ‘mating rate metric’. Simulations with an evolving STI began at the STI-absent host trait values and the results for a reproduction-reducing STI (diamonds) and a viability-reducing STI (triangles) are a three-way evolutionary equilibrium with the STI at evolutionary equilibrium virulence *v* (see Fig. 4). Parameter values (unless shown otherwise): *α* = 10, *μ* = 0.2, *b* = 4, *K* = 1000, *w* = 1, with *c* = 0.0005, *δ* = 0.0005, *d* = 0.02. Each point represents the average of 20 independent simulations +/- standard error. Simulations in which the disease went extinct are excluded from calculating the average. At very low resistance costs, a viability-reducing STI went extinct in 100% of simulations (see Fig. S2) and there is no data for the three-way evolutionary equilibrium.

**Figure S2:**
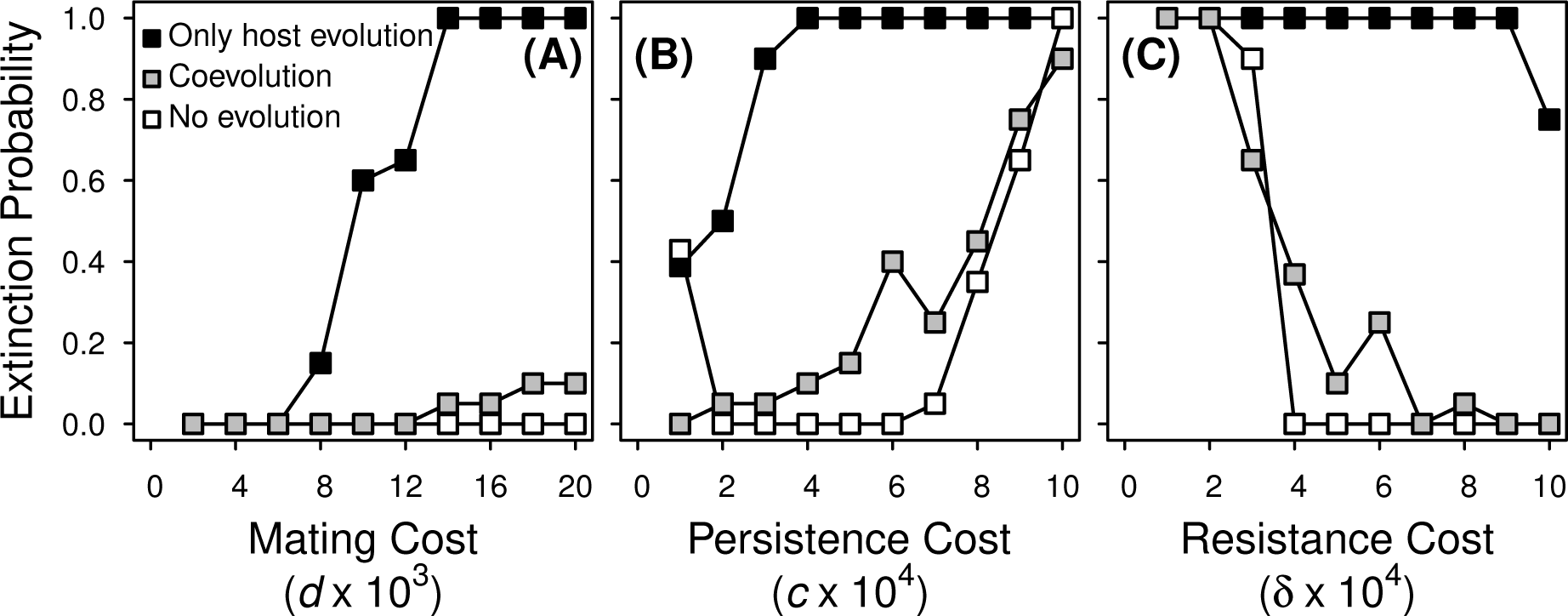
Fraction of simulation runs where a highly virulent viability-reducing STI was driven extinct in hosts paying reproduction costs. The parasite was introduced with *v* = 0.8 at the STI-absent male persistence and female resistance host trait values and neither host nor parasite evolved (open symbols), host traits evolved in the presence of a non-evolving parasite (black symbols), or hosts and STI both evolved (gray symbols). Parameter values (unless shown otherwise): *α* = 10, *μ* = 0.2, *b* = 4, *K* = 1000, *w* = 1, with *c* = 0.0005, *δ* = 0.0005, and *d* = 0.02. The extinction probability was determined from 20 independent simulations.

**Figure S3:**
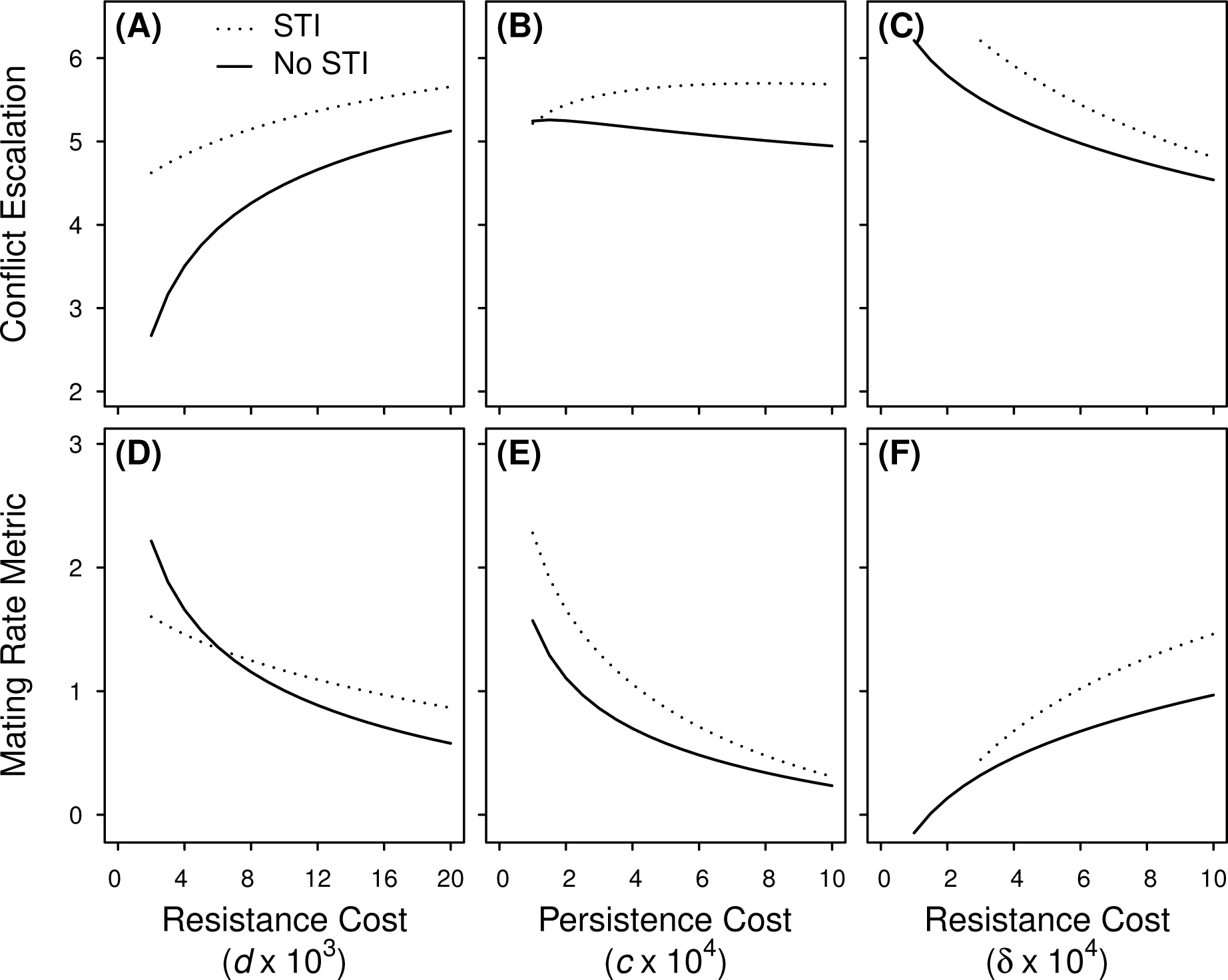
The outcome of coevolution between males and females in the absence of an STI and in the presence of a coevolving viability-reducing STI based on numerical solutions to the analytical model. Both sexes pay viability costs for expressing sexually antagonistic traits. Shown here are the degree of escalation in sexual conflict, measured as the average of *y*\* and *x*\* (*A* - *C*), and the difference between persistence, *y*\*, and resistance, *x*\* (*D* - *F*); values are evolutionarily stable strategies (ESS). The mating rate is an increasing function of the difference between male persistence and female resistance (*y*\* – *x*\*); thus, we label this difference the ‘mating rate metric’. The results shown are a three-way co-ESS with an STI with ESS virulence *v*\* (see Fig. S4). Parameter values (unless shown otherwise): *α* = 0.03, *μ* = 0.2, *b* = 4, *K* = 1000, *w* = 1, with *c* = 0.0005, *δ* = 0.0005, *d* = 0.02. At very low resistance costs, there was no co-ESS with the STI present.

**Figure S4:**
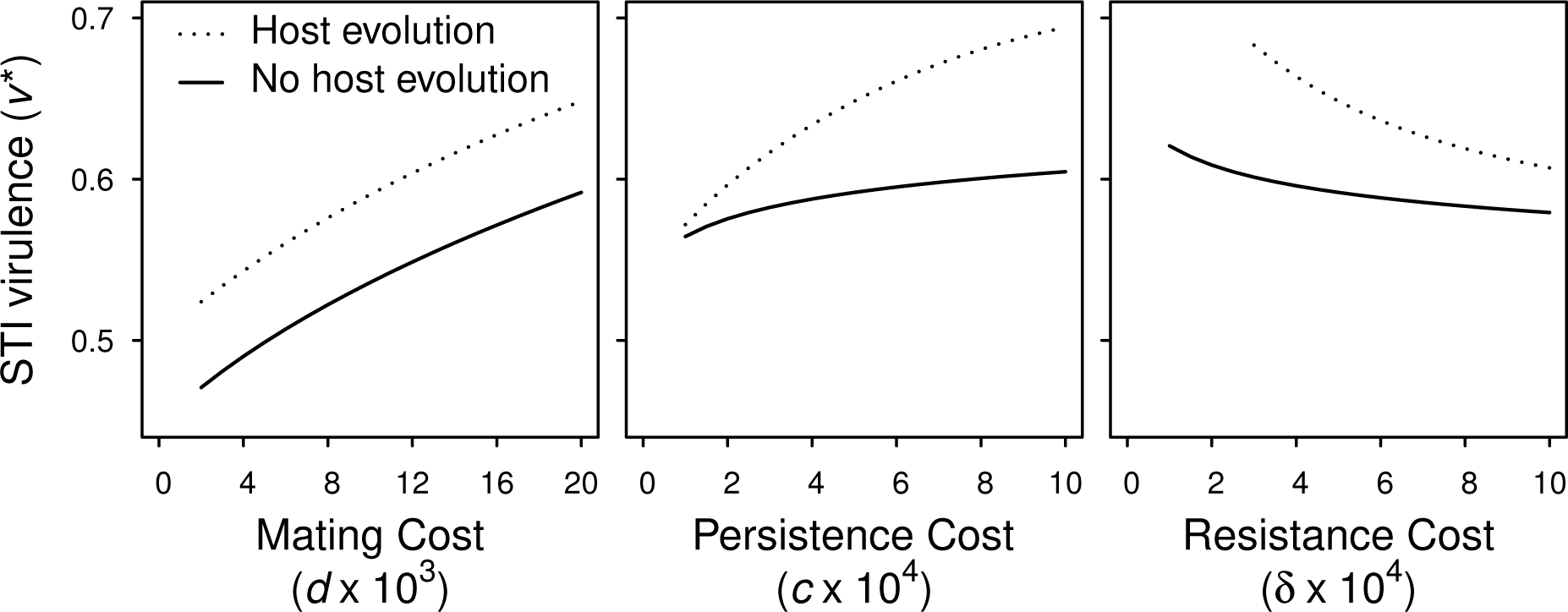
Evolutionary stable virulence (v^*^, ESS virulence) of a viability-reducing STI in a population where hosts do not evolve (open symbols) or evolve (closed symbols). In the absence of host evolution, male persistence and female resistance were treated as constant parameters and set at their STI-absent ESS values. Parameter values (unless shown otherwise): *α* = 0.03, *μ* = 0.2, *b* = 4, *K* = 1000, *w* = 1, with *c* = 0.0005, *δ* = 0.0005, *d* = 0.02. At very low resistance costs, there was no co-ESS with the STI present.

## References

Abbot, P., and L. M. Dill. 2001. Sexually transmitted parasites and sexual selection in the milkweed leaf beetle, Labidomera clivicollis. Oikos 92:91–100.

Alcock, J., C. E. Jones, and S. L. Buchmann. 1977. Male mating strategies in the bee Centris pallida fox (Anthophoridae: Hymenoptera). American Naturalist 111:145–155.

Anderson, R. M., and R. M., May. 1982. Coevolution of hosts and parasites. Parasitology 85:411–426.

Arnqvist, G. 1989. Sexual selection in a water strider: the function, mechanism of selection and heritability of a male grasping apparatus. Oikos 56:344–350.

Arnqvist, G., and Rowe L. 2002. Correlated evolution of male and female morphologies in water striders. Evolution 56:936–947.

Arnqvist, G.. 2005. Sexual Conflict. Princeton University Press, USA.

Ashby, B., and Boots M. 2015. Coevolution of parasite virulence and host mating strategies. Proceedings of the National Academy of Sciences 112:13290–13295.

Bateman, A. J. 1948. lntra-sexual selection in Drosophila. Heredity 2:349–368.

Best, A., and White A. 2009. The implications of coevolutionary dynamics to host-parasite interactions. American Naturalist 173:779–791.

Boots, M., and Knell R. 2002. The evolution of risky behaviour in the presence of a sexually transmitted disease. Proceedings of the Royal Society B: Biological Sciences 269:585–589.

Bro-Jørgensen, J. 2010. Intra- and intersexual conflicts and cooperation in the evolution of mating strategies: lessons learnt From ungulates. Evolutionary Biology 38:28–41.

Day, T., and Burns J. G. 2003. A consideration of patterns of virulence arising from host-parasite coevolution. Evolution 57:671–676.

Ewald, P. W. 1983. Host-parasite relations, vectors, and the evolution of disease severity. Annual Review of Ecology and Systematics 14:465–485.

Fricke, C., Perry, J. T. Chapman, and L. Rowe. 2009. The conditional economics of sexual conflict. Biology Letters 5:671–674.

Gandon, S., P. Agnew, and Y. Michalakis. 2002. Coevolution between parasite virulence and host life-history traits. American Naturalist 160:374–388.

Gavrilets, S., G. Arnqvist, and U. Friberg. 2001. The evolution of female mate choice by sexual conlict. Proceedings of the Royal Society B: Biological Sciences 268:531–539.

Gavrilets, S., and Hayashi T. I. 2006. The dynamics of two-and three-way sexual conlicts over mating. Philosophical Transactions of the Royal Society B: Biological Sciences 361:345–354.

Haddrill, P. R., M. E. N. Majerus, and D. Shuker. 2013. Variation in male and female mating behaviour among different populations of the two-spot ladybird, Adalia bipunctata (Coleoptera: Coccinellidae). European Journal of Entomology 110:87–93.

Immerman, R. S. 1986. Sexually transmitted disease and human evolution: survival of the ugliest. Human Ethology Newsletter 4:6–7.

Immerman, R. S., and Mackey W. C. 1997. Establishing a link between cultural evolution and sexually transmitted diseases. Genetic, Social, and General Psychology Monographs 123:441–460.

Jormalainen, V. 1998. Precopulatory mate guarding in crustaceans: male competitive strategy and intersexual conflict. Quarterly Review of Biology 73:275–304.

Knell, R. 1999. Sexually transmitted disease and parasite-mediated sexual selection. Evolution 53:957–961.

Knell, R. J., and Webberley K. M.. 2004. Sexually transmitted diseases of insects: distribution, evolution, ecology and host behaviour. Biological Reviews 79:557–581.

Kokko, H., Ranta E., G. Ruxton, and P. Lundberg. 2002. Sexually transmitted disease and the evolution of mating systems. Evolution 56:1091–1100.

Lauer, M. J., A. Sih, and J. J. Krupa. 1996. Male density, female density and inter-sexual conflict in a stream-dwelling insect. Animal Behaviour 52:929–939.

Lipsitch, M., and Nowak M. A.. 1995. The evolution of virulence in sexually transmitted HIV/AIDS. Journal of Theoretical Biology 174:427–440.

Lockhart, A. B., P. H. Thrall, and J. Antonovics. 1996. Sexually transmitted diseases in animals: ecological and evolutionary implications. Biological Reviews 71:415–471.

Otto, S. P., and Day T.. 2007. A biologist’s guide to mathematical modeling in ecology and evolution. Princeton University Press, Princeton, NJ.

Rice, W. R., and Holland B.. 1997. The enemies within: intergenomic conflict, interlocus contest evolution (ICE), and the intraspecific Red Queen. Behavioral Ecology and Sociobiology 41:1–10.

Rice, W. R., Stewart, A. D., E. H. Morrow, J. E. Linder, N. Orteiza, and P. G. Byrne. 2006. Assessing sexual conflict in the *Drosophila melanogaster* laboratory model system. Philosophical Transactions of the Royal Society B: Biological Sciences 361:287–299.

Robinson, B. W., and Doyle R. W.. 1985. Trade-off between male reproduction (amplexus) and growth in the amphipod Gammarus lawrencianus. Biological Bulletin 168:482–488.

Rowe, L. 1992. Convenience polyandry in a water strider: foraging conflicts and female control of copulation frequency and guarding duration. Animal Behaviour 44:189–202.

Rowe, L.. 1994. The costs of mating and mate choice in water striders. Animal Behaviour 48:1049–1056.

Rowe, L., G. Arnqvist, A. Sih, and J. Krupa. 1994. Sexual conflict and the evolutionary ecology of mating patterns: water striders as a model system. Trends in Ecology and Evolution 9:289–293.

Rowe, L., E. Cameron, and T. Day. 2005. Escalation, retreat, and female indifference as alternative outcomes of sexually antagonistic coevolution. American Naturalist 165:S5–S18.

Ryder, J. J., K. M. Webberley, M. Boots, and R. J. Knell. 2005. Measuring the transmission dynamics of a sexually transmitted disease. Proceedings of the National Academy of Sciences 102:15140–15143.

Simmons, A. M., and Rogers C. E.. 1994. Effect of an ectoparasitic nematode, Noctuidonema guya-nense, on adult longevity and egg fertility in Spodoptera frugiperda (Lepidoptera: Noctuidae). Biological Control 4:285–289.

Siva Jothy, M. T., and S. J. Plaistow. 1999. A fitness cost of eugregarine parasitism in a damselfly. Ecological Entomology 24:465–470.

Stone, G. N. 1995. Female foraging responses to sexual harassment in the solitary bee Anthophora plumipes. Animal Behaviour 50:405–412.

Thomas, F., E. Oget, P. Gente, D. Desmots, and F. Renaud. 1999. Assortative pairing with respect to parasite load in the beetle *Timarcha maritima* (Chrysomelidae). Journal of Evolutionary Biology 12:385–390.

Thrall, P. H., J. Antonovics, and J. D. Bever. 1997. Sexual transmission of disease and host mating systems: within-season reproductive success. American Naturalist 149:485–506.

Thrall, P. H., J. Antonovics, and A. P. Dobson. 2000. Sexually transmitted diseases in polygynous mating systems: prevalence and impact on reproductive success. Proceedings of the Royal Society B: Biological Sciences 267:1555–1563.

Watson, P. J., G. Arnqvist, and R. R. Stallmann. 1998. Sexual conflict and the energetic costs of mating and mate choice in water striders. American Naturalist 151:46–58.

Webberley, K., G. Hurst, and J. Buszko. 2002. Lack of parasite-mediated sexual selection in a ladybird/sexually transmitted disease system. Animal Behaviour 63:131–141.

